# Validation of deep learning enabled web based and smartphone optimized application RadAnalyzer to measure vertebral heart size and vertebral left atrial size in dogs

**DOI:** 10.1101/2025.11.13.688321

**Authors:** Sonya Gordon, Tomas Reyes, Tabitha Baibos-Reyes, K. Tess Sykes, Alice Watson

## Abstract

**Background:** Objective radiographic measures of heart size including vertebral heart size (VHS) and vertebral left atrial size (VLAS) are associated with inter and intra-observer variability when measured by humans. Artificial intelligence (AI) tools including RadAnalyzer are available to measure VHS and VLAS.

**Objectives:** Compare VHS and VLAS measurements made by web based and smartphone optimized deep learning enabled program, RadAnalyzer, to a trained observer.

**Animals:** 1058 client-owned dogs, across 80 breeds with a variety of heart sizes and thoracic confirmations.

**Methods:** Retrospective, single center, method comparison study. Pearson’s correlation, Bland-Altman plots and Passing-Bablok regression were used to assess agreement.

**Results:** RadAnalyzer measurements of VHS and VLAS correlated well with the human observer’s modified measurements (r=0.917 and r=0.873 respectively) and had small mean biases (0.002 and 0.007 vertebrae respectively).

**Conclusions and clinical importance:** RadAnalyzer had clinically insignificant magnitude differences in measurement of VHS and VLAS when compared to a human observer and can therefore be used to assist veterinarians with measuring VHS and VLAS on right lateral radiographs in dogs of all sizes. Future studies comparing AI derived radiographic measures with echocardiographic measures of cardiac size are required.

## Introduction

Although echocardiography is the gold standard for assessing heart size in dogs [1], echocardiography is not always available. Where echocardiography is not available objective radiographic heart size measurements may be used for staging and monitoring canine cardiac disease [2, 3]. Vertebral heart size (VHS) and vertebral left atrial size (VLAS) have been developed to objectively quantify heart size [4, 5]. Inter and intra-observer variation has been demonstrated for VHS, with documented differences of up to one vertebrae [6, 7] which may have significant effects on sequential measurements on the same dog. Similarly, inter and intra observer variability exists for VLAS [8], and may be associated with clinician experience [9]. Despite these limitations, both VHS and VLAS can be helpful in the clinical setting, especially where echocardiography is not feasible.

Artificial intelligence (AI) is being integrated into various aspects of veterinary medicine, including radiographic and cytological interpretation, as well as predictive models of disease [10, 11]. Within veterinary cardiology AI has been used to detect cardiomegaly in a binary manner [15, 16], as well as to measure VHS [18, 19] and VLAS. A deep learning enabled web based and smartphone optimized application, RadAnalyzer (RadAnalyzer LLC, Austin, USA), was developed to measure VHS and VLAS on lateral thoracic radiographs from dogs. RadAnalyzer is a geometrically informed ensemble of algorithms which was trained on 801 and tested on 199 radiographs. The method used to obtain VHS and VLAS measurements by RadAnalyzer uses an average vertebral size, calculated by taking the mean length of vertebrae four to nine. This modified method positively correlates with the traditional method described by Buchannon and Büchler [5, 12, 13]. Measuring VHS and VLAS with AI has the potential to be a fast, easy, and reliable tool for veterinarians but appropriate validation is essential.

This study aimed to compare the performance of a novel algorithm for measuring VHS and VLAS in dogs, RadAnalyzer, to a trained observer for the measurement of VHS and VLAS on a large dataset of canine right lateral radiographs.

## Materials and Methods

This was a cross-sectional, retrospective, cohort study. Right lateral canine thoracic radiographs obtained from a cohort of cases that were collated for another study were measured by the RadAnalyzer application. The radiographs were originally reviewed prior to inclusion and excluded if there was poor image quality, including over- and under-exposure, or if the required VHS anatomical landmarks (carina, apex, heart base, cranial margin of the heart, caudal vena cava, and the fourth and ninth vertebral body) could not be visualized.

### Manual measurements

A single trained observer measured modified VHS and VLAS (VHS-M and VLAS-M) whereby a mean vertebral length is calculated by measuring from the cranial aspect of T4 to the caudal aspect of T9 and dividing by 5. For VHS the height of the cardiac silhouette was calculated by drawing a line from the center of the ventral aspect of the carina to the ventral most aspect of the cardiac apex and the width of the cardiac silhouette was measured from the ventral aspect of the caudal vena cava to the cranial cardiac waist, perpendicular to the first line. For VLAS a line was drawn from the center of the most ventral aspect of the carina to the most caudal aspect of the left atrium where it intersects with the dorsal border of the caudal vena cava. The number of vertebrae which contribute to the VHS and VLAS were calculated by dividing the sum of the measurements by the mean vertebral length.

### Artificial-Intelligence measurements

RadAnalyzer is a web based and smartphone application. Images of radiographs can be uploaded to the application or photographs of a radiograph are taken within the application which saves and measures the best image and displays the results within 15 seconds. The smartphone application is a unique aspect of this AI radiographic measurement tool. RadAnalyzer assesses images in two stages, stage 1-rough prediction where landmarks are identified and then stage two- a geometrically informed method that fine tunes landmarks identification, images are rejected if prediction is poor. In stage two RadAnalyzer identifies eleven landmarks (carina, cardiac apex, cranial cardiac margin, ventral and dorsal borders of the caudal vena cava as it intersects with the caudal aspect of the cardiac silhouette, the cranial aspect of vertebrae 3, 4 and 5 along and the caudal aspect of vertebrae 8, 9 and 10) for VHS and VLAS calculation (VHS-AI and VLAS-AI).

### Statistical Analysis

Statistical analysis was conducted using R version 4.5.1. Histograms and using Shapiro Wilk tests were used to assess the distribution of continuous data. Correlation between methods was assessed using Spearman correlation. Bland-Altman plots were constructed using the *blandr* package and Passing-Bablok regression using the *mcr* package used to assess agreement between methods. The 95th percentile of absolute difference between VHS-M and VHS-AI were calculated and the 95% confidence intervals computed using bootstrapping resampling with replacement using the *boot* package, the number of bootstrap replicates was set to 10,000. The ground truth was defined as the VHS-M and VLAS-M measurements made by a single, trained observer.

## Results

Radiographs from 1058 dogs representing 106 breeds were included in this study with the most common breeds including: Cavalier King Charles Spaniel (28.4%), Doberman Pinscher (7.7%), Labrador Retriever (7.2%) and Boxer (6.3%) remaining breeds contributed less than 5% each (S1 Table). There was an equal distribution of males (51.0%) and females (49.0%) and the majority were sterilized (83.8%). Median age was 9.4 years (IQR, 6.7-11.2 years, range, 0.4-17.3 years), and median body weight was 11.6 kg (IQR, 7.5-30.4 kg, range, 1.6-90.0 kg). The range of VHS-M and VLAS-M measured by a single trained observer were 8.7-15.5 and 1.0-5.0 vertebrae respectively demonstrating that a wide range of cardiac sizes were included in the study.

VHS was measured on all images by the human and RadAnalyzer. It was not possible for the human observer to measure VLAS in 4 cases and RadAnalyzer algorithm to measure VLAS in 73 different cases, meaning a total of 981 cases had both manual and AI measurements of VLAS.

### Comparison of AI and human

1058 measurements of VHS-M and VHS-AI were available from right lateral radiographic views. For VHS, Pearson’s correlation was 0.917 (95% CI 0.907 to 0.926). Passing-Bablok analysis (Fig 1A) yielded the equation VHS-AI=0.91(VHS-M)+0.97, with 95% CI of 0.89-0.93 for the slope and 0.73 to 1.21 for the intercept. Bland–Altman analysis (Fig 1B) showed a bias of 0.002 (SE 0.013; 95% CI −0.024 to 0.029) between VHS-M and VHS-AI. The 95^th^ percentile absolute difference between VHS-M and VHS-AI was 0.88 vertebrae (95% CI 0.83-0.97, Fig 2).

**Fig 1:**
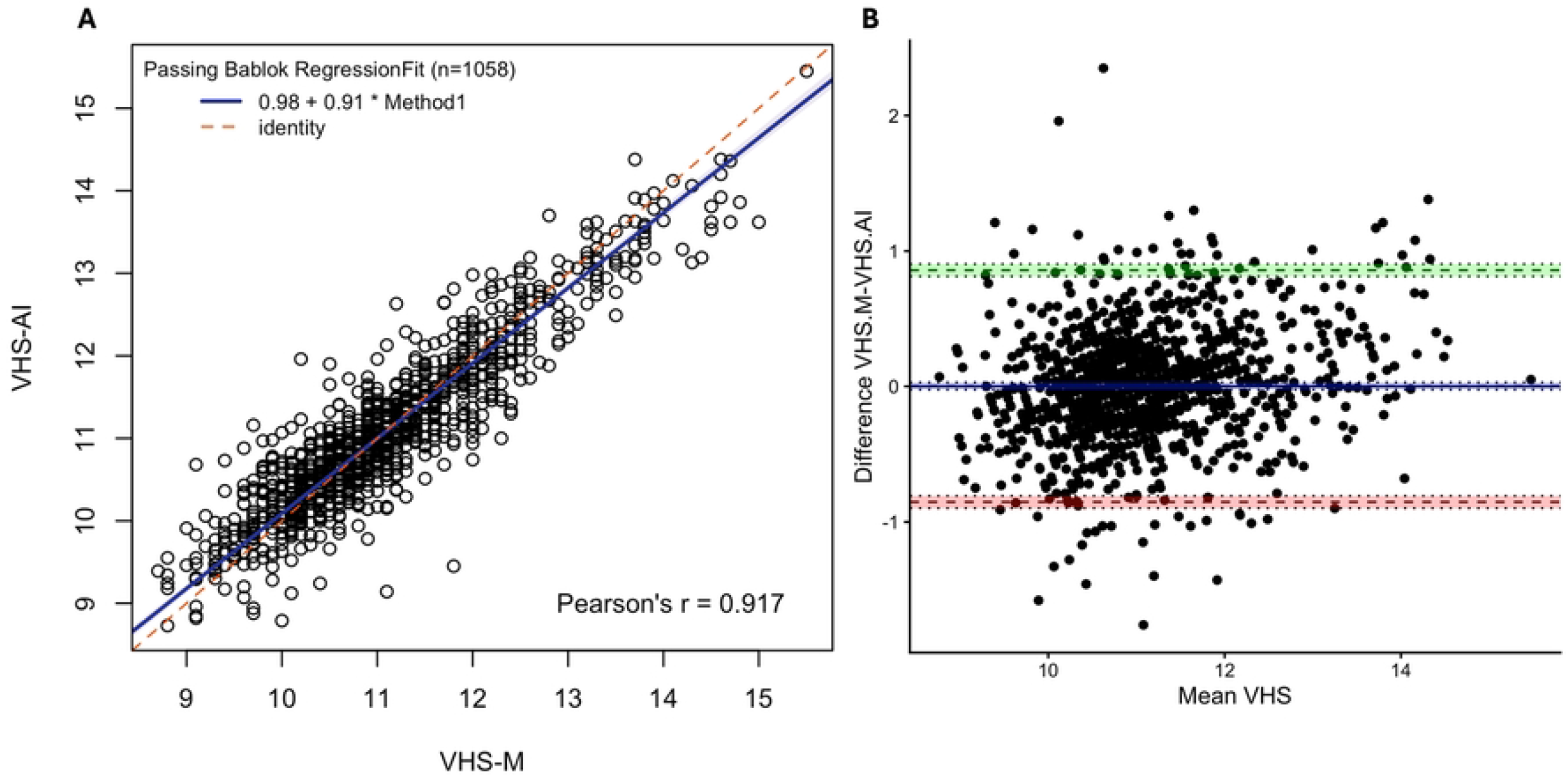
Comparison of VHS measurement by human and RadAnalyzer. Passing– Bablok analysis (A) analysis yielded the equation VHS-AI=0.91(VHS-M)+0.98, Pearson’s linear correlation coefficient r=0.917, n=1058. Blue line: fitted regression line, shading: 95% confidence interval, red dashed line: identity. Bland–Altman analysis (B) demonstrated a mean bias of 0.002 vertebrae (95% CI −0.023 to 0.029) for the VHS-AI method. Dashed lines give limits of agreement (±1.96 SD) and mean bias; shading gives 95% confidence intervals with dotted lines at the limits. VHS indicates vertebral heart size; VHS-AI, VHS measured by RadAnalyzer, VHS-M, VHS measured by a human observer and SD, standard deviation.

**Fig 2:**
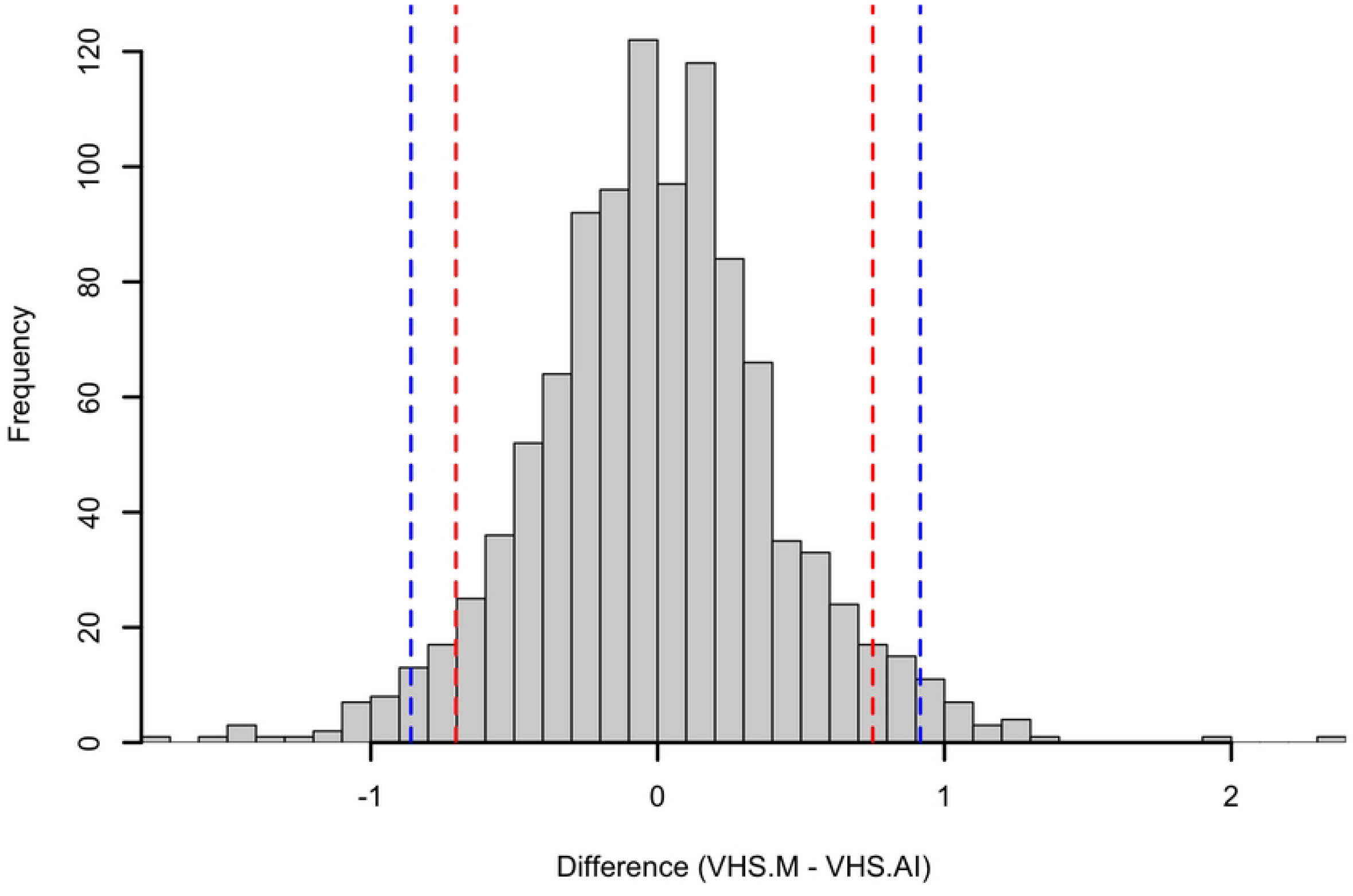
Histogram of the difference between modified vertebral heart size (VHS) determined by human observer (VHS.M) and Radanalyzer (VHS.AI). The red lines indicate the 5th percentile and 95th percentile. The blue dashed lines represent the 2.5th percentile and the 97.5th percentile.

981 measurements of VLAS-M and VLAS-AI were available from right lateral radiographic views and Pearson’s correlation was 0.873 (95% CI 0.857 to 0.887). Passing-Bablok analysis (Fig 3A) yielded the equation VLAS-AI=0.90(VLAS-M)+0.20, with 95% CI of 0.88-0.93 for the slope and 0.13 to 0.25 for the intercept. Bland–Altman analysis (Fig 3B) showed a bias of 0.007 (SE 0.007; 95% CI −0.004 to 0.025) between VLAS-AI and VLAS-M. The 95^th^ percentile absolute difference between VLAS-M and VLAS-AI was 0.49 vertebrae (95% CI 0.44-0.51, Fig 4).

**Fig 3:**
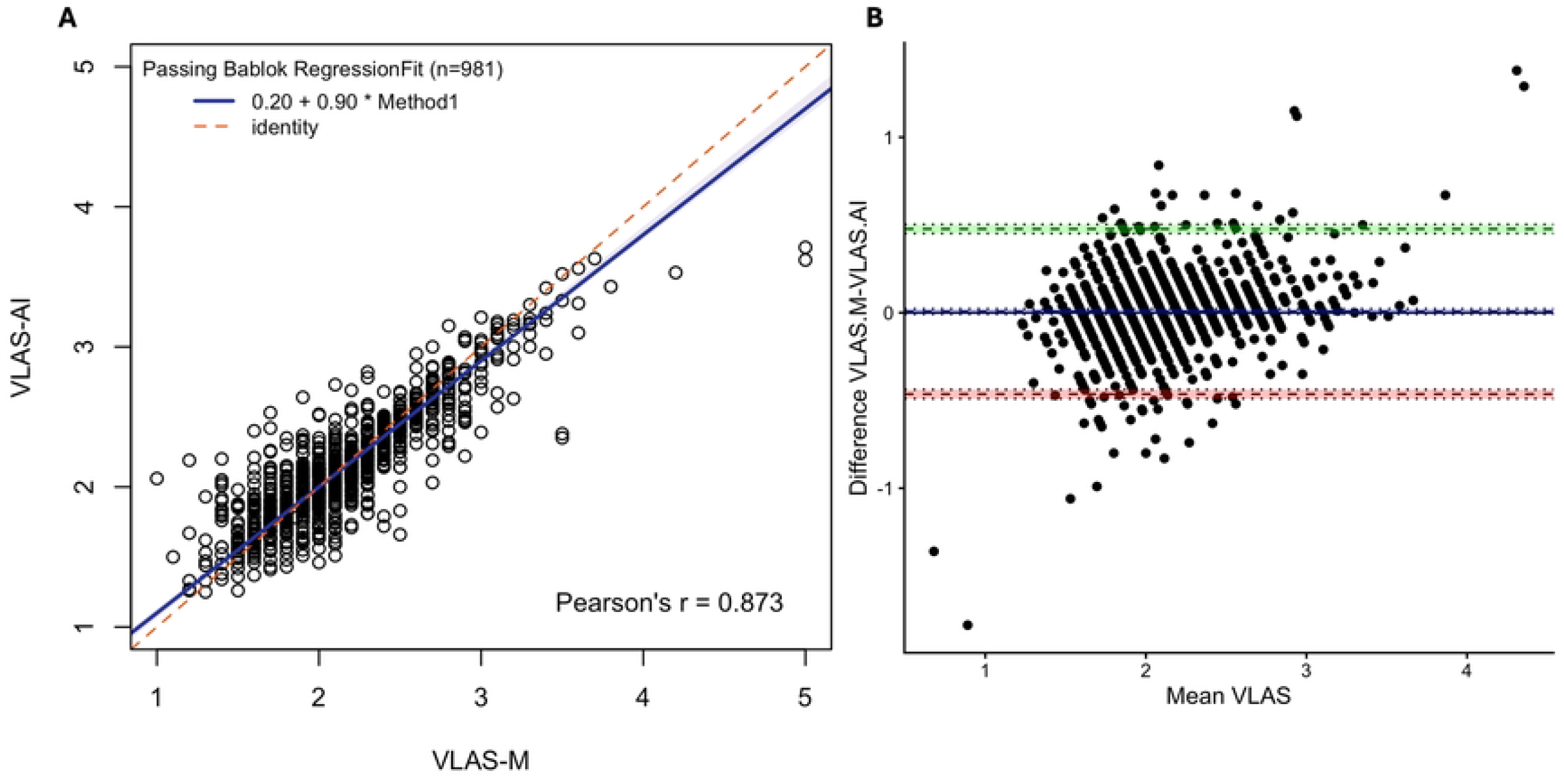
Comparison of VLAS measurement by human and RadAnalyzer. Passing– Bablok analysis. (A) yielded the equation VLAS-AI=0.90(VLAS-M)+0.20, Pearson’s linear correlation coefficient r=0.873, n=981. Blue line: fitted regression line, shading: 95% confidence interval, red dashed line: identity. Bland–Altman analysis (B) demonstrated a mean bias of 0.01 vertebrae (95% CI −0.003 to 0.03) for the VHS-AI method. Dashed lines give limits of agreement (±1.96 SD) and mean bias; shading gives 95% confidence intervals with dotted lines at the limits. VLAS indicates vertebral left atrial size; VLAS-AI indicates VLAS measured by RadAnalyzer, VLAS-M, VLAS measured by a human observer and SD, standard deviation.

**Fig 4:**
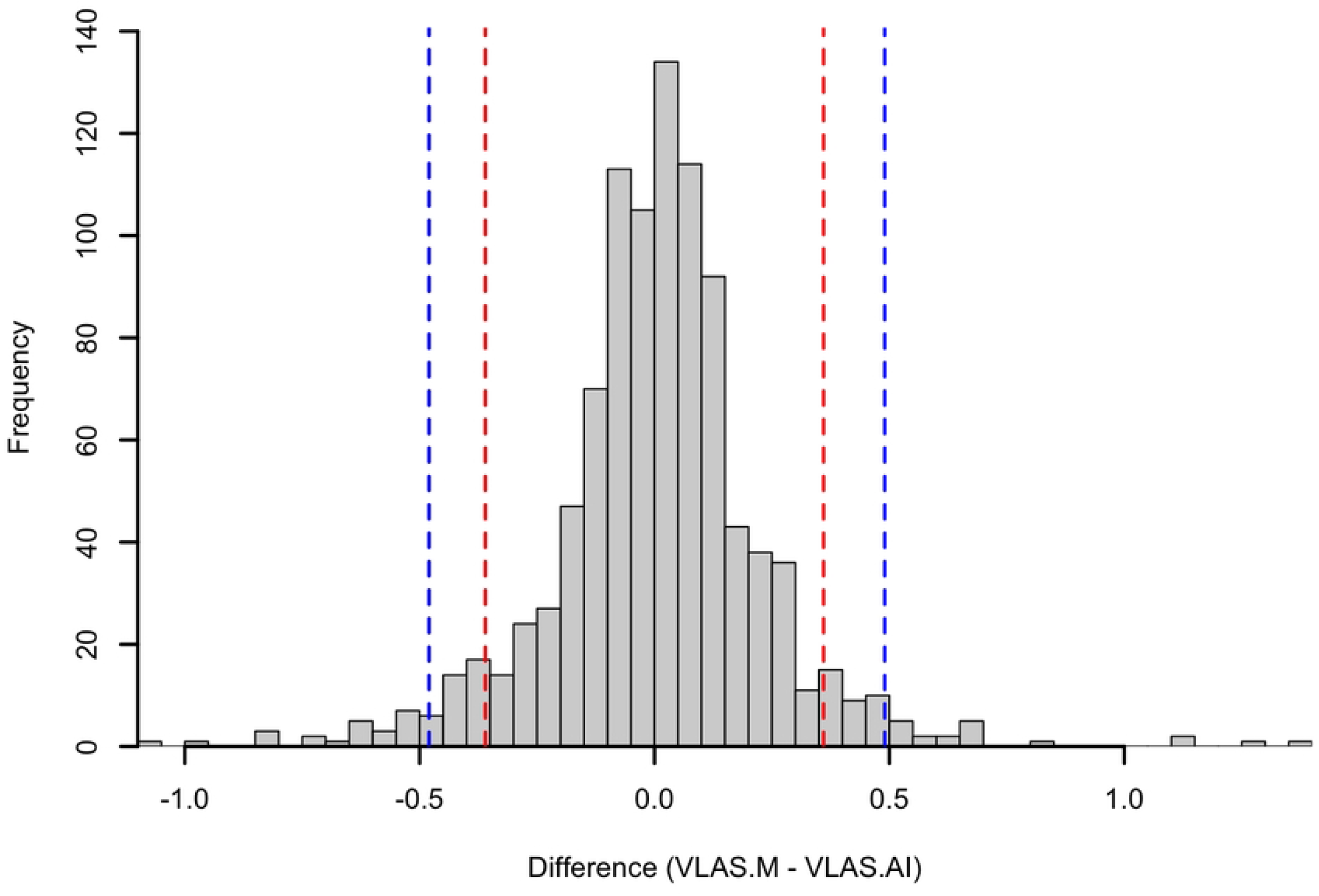
Histogram of the difference between modified vertebral left atrial size (VLAS) determined by human observer (VLAS.M) and Radanalyzer (VLAS.AI). The red lines indicate the 5th percentile and 95th percentile. The blue dashed lines represent the 2.5th percentile and the 97.5th percentile.

## Discussion

Deep-learning enabled software, RadAnalyzer, correlated better with trained observer for measuring VHS (r=0.917, 95% CI 0.907 to 0.926) than VLAS (r=0.873 (95% CI 0.857 to 0.887). Minimal bias was present for both VHS (0.002, 95% CI −0.024 to 0.029) and VLAS (0.007, 95% CI −0.004 to 0.025). Differences between RadAnalyzer and a single human observer for both VHS and VLAS were minimal and are unlikely to be of a clinically relevant magnitude, suggesting that RadAnalyzer may be useful in clinical practice. RadAnalyzer may reduce interobserver variability associated with VHS and VLAS measurements.

The mean bias derived from Bland-Altman plots between AI and human for VHS and VLAS in this study was 0.001 and 0.007 vertebrae respectively which are unlikely to cause clinically significant differences in measurements. The mean bias for VHS is lower than a previous study using different AI-algorithms to measure VHS where measurements differed by 0.09 vertebrae to board-certified radiologists [18]. Both AI algorithms give less variation than the 1.05 vertebrae that was observed between 16 human observers in another study [6]. RadAnalyzer can be deployed outside of a picture archiving and communication system (PACS) system both on a computer and by using the RadAnalyzer app on a smart phone to capture a picture of a radiograph from any screen. VHS and VLAS are time-consuming methods for assessing cardiac and left atrial sizes in dogs which are subject to interobserver variability, with more experienced individuals typically performing better [6, 9, 14]. The goal of AI in veterinary medicine should be to empower veterinarians, not replace them. AI can provide reliable information to help veterinarians make informed decisions, but it should not be expected to replace their expertise. However, AI can improve efficiency and precision and therefore allow veterinarians’ time to focus on more complex and high-value tasks, such as diagnosing and treating patients.

One of the key challenges in AI is explainability. Explainable AI aims to provide transparency into how the AI came up with an answer, but the truth is that many AI systems, including deep learning frameworks, are still black boxes. Although this algorithm can offer some understanding of its decisions for VHS and VLAS, deep learning systems remain complex and difficult to interpret—a common challenge in AI technology. In the RadAnalyzer system the landmarks required for a human to measure VHS and VLAS using both the traditional and modified methods are shown on the digital image, enabling veterinarians to check and confirm landmark selection and the measurements generated by the AI algorithm and if judged to be incorrect the annotated landmarks can be edited and the VHS/VLAS is immediately recalculated. A benefit of RadAnalyzer is that it will not measure VHS or VLAS on radiographs considered to be non-diagnostic, reducing overinterpretation of poor-quality radiographs which may cause erroneous conclusions to be drawn and potentially affecting patient care.

If echocardiography is unavailable, radiographic indices of heart size, including VHS and VLAS, can be utilized for the diagnosis, staging and monitoring of canine cardiac disease [2, 3, 15]. This study was conducted because echocardiography is not readily available to many practicing veterinarians and there is variability between individuals when measuring VHS and VLAS [6, 14]. Deep-learning technology eliminates inter-observer and intra-observer variability associated with human observers [6, 14, 16]. Reliable measurements and monitoring cardiac size is important as the rate of increase in VHS accelerates just before the onset of CHF in myxomatous mitral valve disease and a rate of change above >0.08 vertebral units per month is associated with the onset of CHF [17, 18]. Similarly, a rate of change of increase in VLAS above 0.02 per month is associated with the onset of CHF in myxomatous mitral valve disease [19]. RadAnalyzer may therefore be particularly useful for repeated measurements and longitudinal monitoring.

There are several limitations to this study. A single observer obtained the manual VHS and VLAS measurements (VHS-M and VLAS-M), to reduce the impact of interobserver variability [6], intraobserver variability was not evaluated. This observer was also considered the ground truth, despite it being accepted that humans are inherently variable [6, 12]. Future studies could compare agreement between radiographic measurements from RadAnalyzer with echocardiographic measurements of cardiac size as echocardiography is considered the most highly sensitive and specific method for the diagnosis and staging of canine cardiac disease and for measurements of cardiac size. This study also did not include patients with active heart failure, or other pulmonary infiltrate patterns as pulmonary opacities may affect performance. The effect of radiographic pulmonary infiltrates on the performance of the RadAnalyzer algorithm and use of RadAnalyzer in different clinical settings and populations could be investigated in future studies. Radanalyzer cannot distinguish between right and left lateral radiographs and has not been validated on left lateral radiographs.

## Conclusion

The commercially available web based and smartphone application, RadAnalyzer, generates comparable measurements of VHS and VLAS to a trained veterinarian and will reduce the impact of interobserver variability. Following AI annotation and measurement, it is possible to view landmarks, enabling clinicians to confirm the validity or adjust as appropriate of the annotated landmarks associated with the measurement to avoid mistakes in clinical decision making. Further studies are needed to compare agreement between measurements from this tool with echocardiographic variables, which may further support the use of this tool in making clinical decisions for patients with cardiac disease.

## Acknowledgments

The authors thank the cardiology faculty, residents and technicians at the Texas A&M University, College of Veterinary Medicine and Biomedical Sciences for their assistance with data collection and acknowledge the support of the Aggies Invent innovative design competition.

## Supporting information captions

**S1 Table. Dog breeds**. Number and percentage of dogs of each breed included within the study.

